# Bat eye movements resolve a long-standing question in gaze control

**DOI:** 10.64898/2026.03.10.710949

**Authors:** Hui Ho Vanessa Chang, Grace Capshaw, Dimitri Skandalis, Cynthia F. Moss, Kathleen E. Cullen

**Author notes:** **Author contact:** Hui Ho Vanessa Chang Grace Capshaw, Dimitri Skandalis Cynthia F. Moss. **Lead contact** Kathleen E. Cullen.

## Abstract

Eye movements enable visual information gathering and stabilize gaze via optokinetic (OKR) and vestibulo-ocular reflex (VOR) pathways.^1^ Echolocating bats, despite their rapid and agile flight maneuvers to land upside down and navigate 3D space, have long been thought not to move their eyes, an assumption originating from Walls’s influential assertion over 80 years ago^2^ but never tested with empirical measurements. Here we present quantitative analysis of eye movements driven by visual and vestibular signals in Seba’s short-tailed bat (*Carollia perspicillata*). Bats generated robust visually driven OKR with an oculomotor range of ∼±10°, and displayed strong otolith-mediated responses during off-vertical axis rotation. In contrast, they showed minimal semicircular canal–driven angular VOR (aVOR) for passive head rotations that elicit large, sustained responses in mice. Micro-CT reconstructions revealed that bats and mice have similar semicircular canal geometry, indicating that the weak aVOR does not reflect peripheral anatomical constraints. These findings provide the first empirical demonstration that bats make robust eye movements and exhibit strong visual and otolith-driven components of gaze stabilization. We propose that semicircular canal signals may be more strongly engaged during active flight and modulated by behavioral state–dependent tuning of vestibular pathways to support ecologically specialized behaviors.

## Results

### Bats Generate Robust Optokinetic Eye Movements

We first compared visually driven eye movements of bats with those of mice, which have been relatively well studied (e.g.,^3,4^) and are comparable in size and weight. To evoke optokinetic eye movements, animals were head-fixed and placed at the center of a visual surround rotating about the earth-vertical axis at a constant velocity (Figure 1A; see Methods). As expected, mice exhibited robust compensatory visual tracking, characterized by large-amplitude slow-phase eye movements with amplitudes approaching ±20° and reliable quick-phase resets (Figures 1B and 1C, blue traces). These visually driven responses were sustained across cycles and showed consistent tracking across both stimulus directions. (Figures 1C and 1D, blue traces).

**Figure 1.**
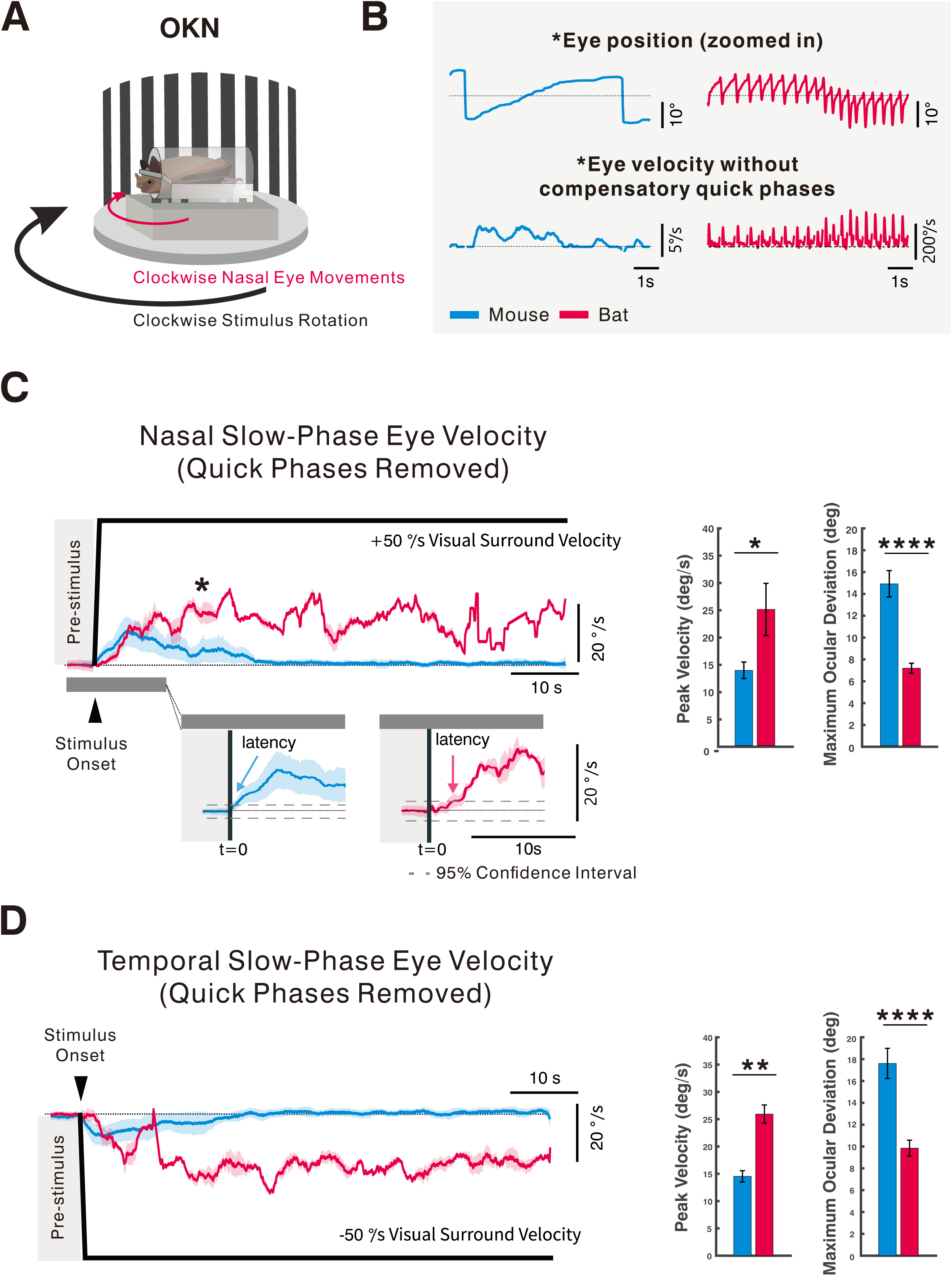
Seba’s short-tailed bat exhibited robust OKR responses to constant velocity stimulation. (**A**) Experimental setup for recording optokinetic reflex (OKR) eye movements during constant-velocity rotation of the visual surround. (**B**) Selected averaged eye-position traces from bats (red) and mice (blue), showing alternating slow-phase tracking and quick-phase resets. Traces are extracted from the time window indicated by the asterisk. In bats, individual slow phases often comprised a transient, higher-velocity eye movement followed by a slower tracking component, with both components occurring in the direction of the visual stimulus (see also Figure S1). (**C**) Nasal-direction OKR evoked by +50°/s (clockwise) visual surround rotation. Traces are time-aligned to stimulus onset (t=0), with a pre-stimulus baseline epoch from-4 to 0 s. *Top*: Average slow-phase eye velocity (mean ± SEM) for bats (N = 4) and mice (N = 6) during +50°/s (clockwise) visual surround rotation, which evokes nasal eye movements. Mean slow-phase eye velocity was computed by pooling valid slow-phase segments across animals. The black trace indicates surround velocity. Mice reached peak nasal slow-phase velocities of 13.7 ± 1.5°/s, whereas bats exhibited significantly higher peak velocities (24.7 ± 4.7°/s; p = 0.038, Mann–Whitney U test). Maximum ocular deviation—defined as the largest slow-phase eye-position displacement from baseline during the stimulus—was 14.7 ± 1.2° in mice and 7.0 ± 0.5° in bats (p < 0.0001). *Bottom*: Zoomed view of the shaded region showing pre-and post-stimulus intervals. Dashed horizontal lines denote the 95% confidence interval (CI) of baseline slow-phase eye velocity. Arrows indicate response latency, defined as the first time point at which the 95% CI of the mean post-stimulus response exceeded the baseline CI, demonstrating a statistically reliable OKR response. Insets summarize mean peak velocity (left) and maximum ocular deviation (right). Latency estimates were 1.4 s for mice and 3.7 s for bats using this criterion. (**D**) Temporal-direction OKR evoked by −50°/s (counterclockwise) visual surround rotation, plotted with the same conventions as in (**C**). Mice reached peak temporal velocities of 14.5 ± 1.0°/s, whereas bats exhibited significantly higher temporal peak velocities (25.9 ± 1.7°/s; p = 0.0095). Maximum ocular deviation was 9.8 ± 0.7° in bats and 17.6 ± 1.4° in mice (p < 0.0001). Latencies for temporal OKR were 0.4 s in mice and 2.4 s in bats, based on the same baseline CI-crossing criterion. All data are shown as mean ± SEM. *P ≤ 0.05; **P ≤ 0.01; ****P ≤ 0.0001. See also Figure S1.

Quantitative OKR metrics for mice, nasal–temporal and temporal–nasal peak velocities, maximum ocular deviation, and response latency estimates, are reported in the Figure 1 legend.

Seba’s short-tailed bats (*Carollia perspicillata*) also, unexpectedly, generated a clear and well-structured OKR (Figures 1B and 1C, red traces). As in mice, bat OKR responses comprised alternating slow-phase eye movements that tracked the stimulus and oppositely directed quick-phase resets that recentered the eyes within the oculomotor range. However, in bats the slow phase itself was not temporally uniform. Inspection of individual slow phases revealed a two-component structure, with both components occurring in the direction of the visual stimulus: an initial, brief higher-velocity transient followed by a slower, sustained tracking component (Figure 1B). This two-component structure was consistently observed across individual bats (Figure S1). As a consequence, when slow-phase responses were pooled across trials and animals, variability in the timing and magnitude of this initial transient contributed to the increased dispersion observed in the averaged slow-phase velocity traces (Figures 1C and 1D). Despite this variability, OKR amplitudes in bats approached ±10°, and bats exhibited significantly larger peak slow-phase velocities than mice across both stimulus directions (Figure 1C). Detailed quantitative metrics for both species, including nasal–temporal and temporal–nasal peak velocities, maximum ocular deviation, and latency estimates, are provided in the Figure 1 legend.

Together, these findings establish that *Carollia perspicillata* generates robust visually driven eye movements. The presence of a well-structured OKR indicates that bats use eye movements for visual tracking and that their visuomotor pathways are active under passive visual stimulation.

### Semicircular Canal–Driven aVOR Is Minimal in Bats

We next examined vestibularly driven eye movements by quantifying the angular vestibulo-ocular reflex (aVOR) during passive whole-body rotation in darkness (Figure 2A), a classic paradigm for isolating semicircular canal–driven gaze stabilization. As expected, mice exhibited a robust aVOR response (Figures 2B and 2C, blue traces), with compensatory slow-phase eye movements that closely matched head velocity and directionally appropriate quick phases that maintained gaze within the oculomotor operating range. Over time, slow-phase eye velocity decayed gradually, consistent with engagement of the velocity-storage mechanism, a hallmark of mammalian aVOR dynamics. Overall, these responses are consistent with the well-described canal-driven stabilization observed in rodents.

**Figure 2.**
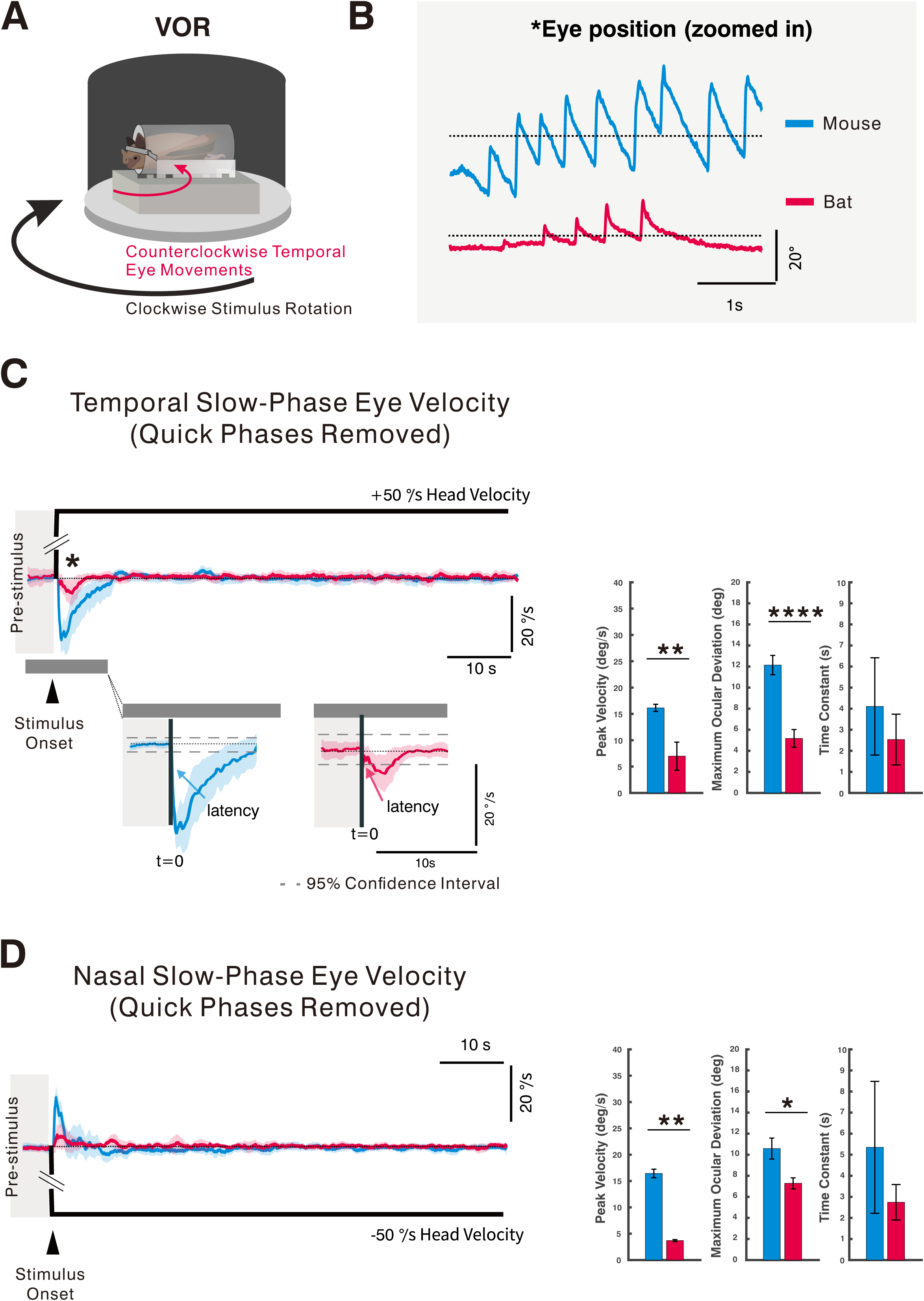
Semicircular canal–driven vestibulo-ocular reflex (aVOR) is minimal in Seba’s short-tailed bats. (**A**) Experimental setup for measuring the angular vestibulo-ocular reflex (aVOR) during passive constant-velocity rotation of the turntable in darkness. (**B**) Example eye-position traces from bats (red) and mice (blue), showing extracted slow and quick phases during the analysis window indicated by the asterisk. (**C**) *Top*: Average slow-phase eye velocity of bats (N = 4) and mice (N = 6) in response to +50°/s (clockwise) turntable rotation, which elicits temporal compensatory eye movements. Traces are time-aligned to stimulus onset (t=0), with a pre-stimulus baseline epoch from-4 to 0 s. The black trace indicates table velocity. In mice, constant-velocity rotation evoked a strong compensatory response that decayed exponentially, consistent with velocity-storage dynamics (time constant: 4.11 ± 2.3 s), with a peak temporal eye velocity of 16.1 ± 0.7°/s. In contrast, bats generated only a brief onset transient with a significantly smaller peak velocity (6.9 ± 2.7°/s; p = 0.0095, Mann–Whitney U test) and reduced maximum ocular deviation (5.2 ± 0.8°; p < 0.0001). The temporal decay time constant in bats was 2.5 ± 1.2 s. *Bottom:* Zoom-in of the shaded region highlighting pre-and post-stimulus epochs. Dashed lines indicate the 95% confidence interval (CI) of the pre-stimulus baseline. Arrows denote response latency, defined as the first time point at which the mean slow-phase eye velocity exceeded the baseline 95% CI, confirming responses exceeded noise levels (0.2 s in mice; 0.1 s in bats). Insets show mean peak velocity (left), maximum ocular deviation (middle), and decay time constant (right). (D) Average slow-phase eye velocity during −50°/s (counterclockwise) rotation, eliciting nasal compensatory eye movements, with conventions as in (C). Mice reached a peak nasal eye velocity of 16.4 ± 0.8°/s with a decay time constant of 5.3 ± 3.1 s and a latency of 0.039 s. Bats showed markedly reduced nasal responses, with a peak eye velocity of 3.7 ± 0.2°/s (p = 0.0095) and maximum ocular deviation of 7.3 ± 0.5° (p = 0.0106). The nasal decay time constant in bats was 2.7 ± 0.8 s. Latencies were 0.2 s in mice and 0.6 s in bats using the same CI-crossing criterion. Insets show mean peak velocity (left), maximum ocular deviation (middle), and decay time constant (right). Data are mean ± SEM. *P ≤ 0.05; **P ≤ 0.01; ****P ≤ 0.0001.

In striking contrast, Seba’s short-tailed bats (*C. perspicillata*; red traces) exhibited a substantially smaller, transient eye movement at motion onset in response to the same rotational stimulus (Figures 2B and 2C), which was consistent across individuals. Compared to mice, peak eye velocities were markedly reduced and decayed rapidly, resulting in little sustained compensatory tracking. Quantitative comparisons revealed a clear species difference: mice generated significantly larger, compensatory eye velocities, whereas bats showed substantially smaller aVOR responses (compare Figures 2C and 2D). Corresponding quantitative metrics for both species—including peak velocities, maximum ocular deviation, and decay time constants—are provided in the Figure 2 legend.

Together, these findings demonstrate that *C. perspicillata* exhibits little to no canal-driven aVOR under passive rotational stimulation. The presence of only a brief onset transient indicates that the semicircular canals are functional, but that canal signals do not drive sustained oculomotor output during passive body rotation.

### Otolith-Driven Responses Are Strong and Reliable in Bats

We next isolated otolith contributions using off-vertical axis rotation (OVAR), which elicits a well-defined steady-state modulation of eye velocity aligned with the changing orientation of the gravito-inertial vector (Figure 3A). As expected, mice (blue traces) displayed the classical two-component response: an initial canal-driven transient followed by a strong sinusoidal modulation during constant-velocity rotation that reflects otolith encoding of tilt (Figures 3B and 3C).

**Figure 3.**
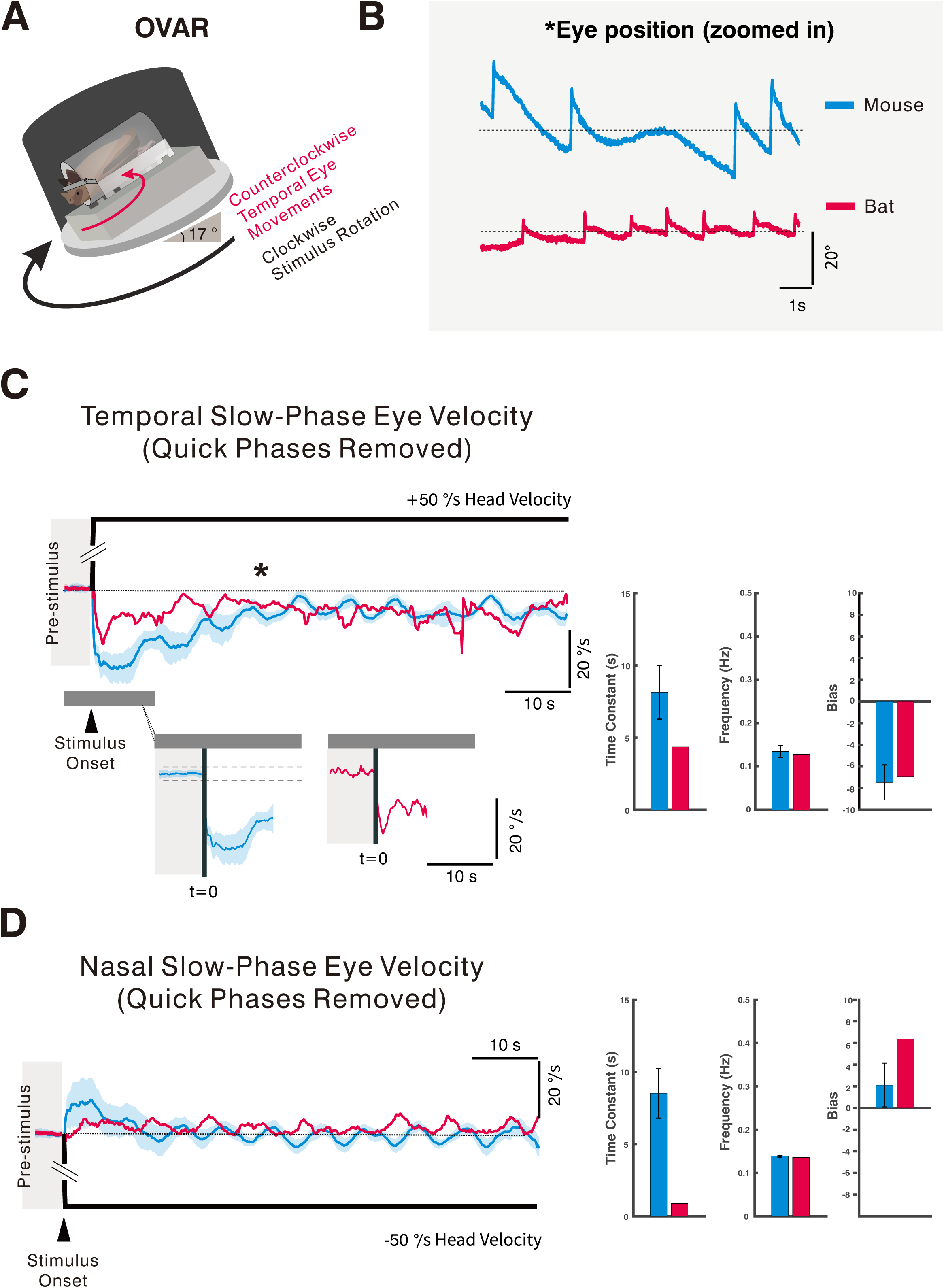
Seba’s short-tailed bats exhibit robust otolith-driven responses during off-vertical axis rotation (**A**) Experimental setup for acquiring otolith-driven responses using off-vertical axis rotation (OVAR) with the turntable tilted 17° from earth vertical. (**B**) Example eye-position traces showing quick and slow phases for mice (blue) and a bat (red), extracted from the window indicated by the asterisk. (**C**) *Top:* Average slow-phase eye velocity of mice (N = 6) and a bat during +50°/s (clockwise) OVAR, which elicits temporal eye movements. Traces are time-aligned to stimulus onset (t=0), with a pre-stimulus baseline epoch from-4 to 0 s. The black trace indicates turntable velocity. Mice exhibited an initial canal-mediated transient followed by a steady-state sinusoidal modulation reflecting otolith-driven encoding of gravito-inertial tilt. The oscillation frequency was 0.13 ± 0.01 Hz in mice and 0.13 Hz in the bat. The bias component (mean velocity offset) and peak temporal velocities were comparable across species. *Bottom:* Zoom-in of the shaded region highlighting pre-and post-stimulus epochs. Dashed lines denote the 95% confidence interval of the pre-stimulus baseline. Insets show mean peak velocity (left), oscillation frequency (middle), and bias (right). (**D**) Average slow-phase eye velocity for −50°/s (counterclockwise) rotation, eliciting nasal eye movements, plotted using the same conventions as in (**C**). As for temporal rotation, mice displayed a canal-mediated transient followed by a sinusoidal otolith-driven steady-state response. The nasal oscillation frequency was 0.14 ± 0.00 Hz in mice and 0.14 Hz in the bat. Peak nasal velocities and bias were comparable between species, and the bat exhibited a robust steady-state OVAR response similar to that of mice. Insets show mean peak velocity (left), oscillation frequency (middle), and bias (right). Data are presented as mean ± SEM.

Quantitative measures of response time constant, frequency, and bias are provided in the Fig. 3 legend.

Remarkably, Seba’s short-tailed bats (*C. perspicillata*; red traces) also exhibited a robust sinusoidal steady-state modulation during OVAR, comparable in amplitude to that of mice (Figures 3B and 3C). Despite the minimal aVOR response under purely rotational stimulation, the bat generated a clear and consistent otolith-driven signal under OVAR. Quantification revealed that both peak eye velocity and modulation amplitude in bats closely matched those of mice for OVAR applied in either direction (Figures 3C and 3D). Population-level comparisons confirmed that otolith-mediated responses were strong in both species, with no significant difference in modulation amplitude (Figures 3C and 3D, compare blue and red bars, respectively). Corresponding fit parameters are reported in the Figure 3 legend.

Together, these findings demonstrate that otolith pathways are robustly preserved in *C. perspicillata* and contribute robustly to gaze stabilization. The dissociation between canal-driven and otolith-driven responses indicates that bats selectively engage vestibular modalities and place substantial weight on gravito-inertial cues, even under passive motion.

### Bats Lack Frequency-Dependent Modulation in Visual, Vestibular, and Combined Stimuli

To further examine sensory weighting in bats, we quantified frequency responses of the OKR, angular VOR in dark (VOR_d_), and visually enhanced VOR (VOR). In mice (blue traces), VOR_d_ gain increased systematically with stimulus frequency, approaching unity at higher frequencies, whereas OKR gain declined, yielding the classical OKR–VOR crossover (Figures 4A and 4B). When visual and vestibular stimuli were combined, mice exhibited clear multisensory enhancement, producing higher gains than either input alone (Figure S2). Gain–frequency relationships for both OKR and VOR_d_ in mice are provided in the Figure 4 legend. Bats (*C. perspicillata*; red traces), however, showed no such frequency dependence, with VOR_d_ gain near zero across all frequencies tested, consistent with their lack of canal-driven eye stabilization. In addition, bat OKR gain did not decrease at higher stimulation frequencies; instead, it remained uniformly low across the tested range (Figures 4A and 4B). OKR gains in bats were uniformly low and remained relatively flat, indicating that bat OKR dynamics markedly differ qualitatively from the frequency-dependent behavior observed in mice. Further, multisensory stimulation during VOR failed to enhance performance in bats: VOR responses were indistinguishable from OKR across the entire frequency range (Figure S1). Corresponding gain–frequency curves for bats are likewise reported in the Figure 4 legend.

**Figure 4.**
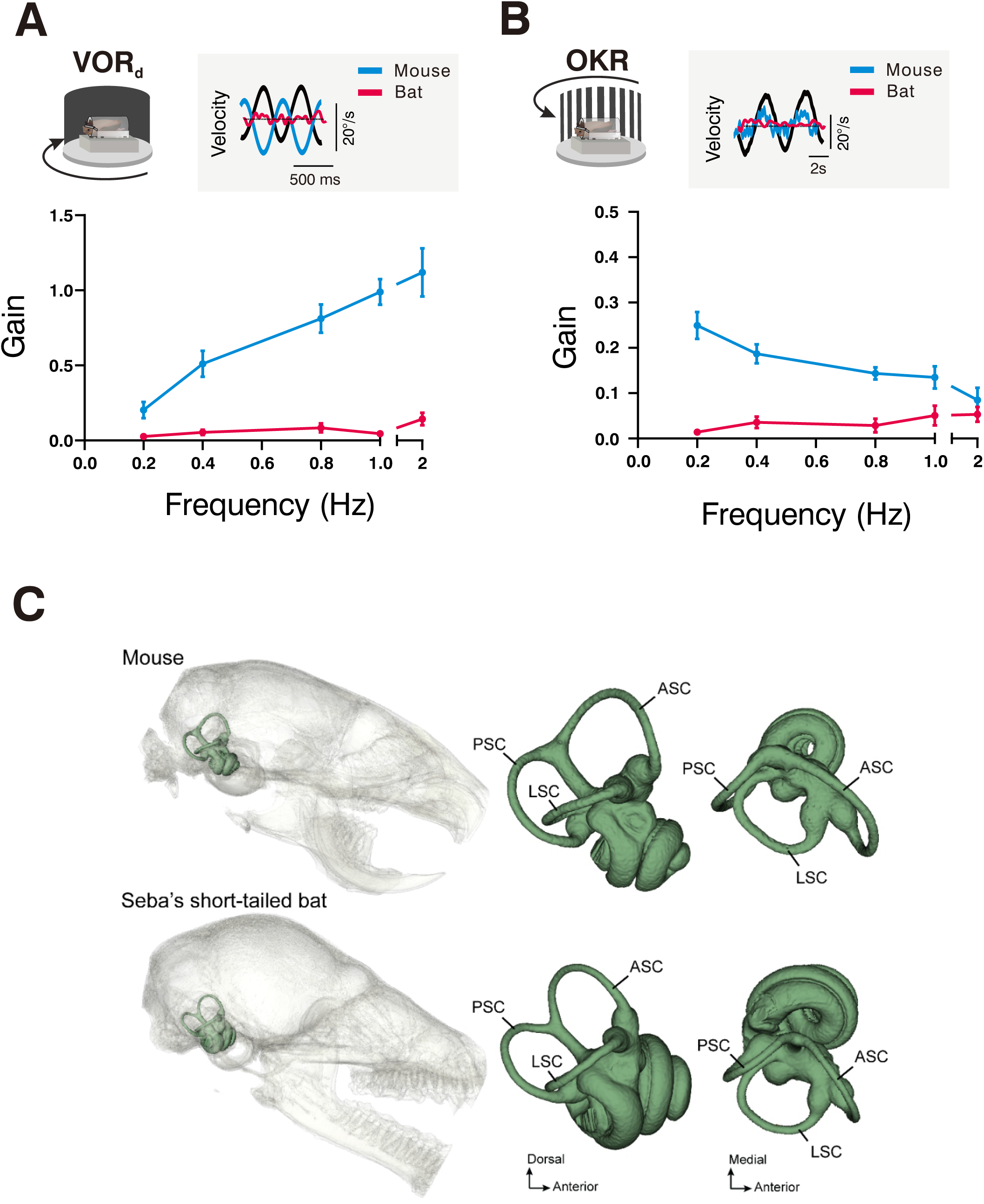
Bats lack frequency-dependent gain modulation despite preserved inner-ear morphology (**A**) Angular vestibulo-ocular reflex in dark (VOR_d_) gain plotted as a function of stimulus frequency for control mice (blue; N = 6) and Seba’s short-tailed bats (red; N = 4). Data are mean ± SEM. The inset shows example eye-velocity traces at 2 Hz for mice (blue) and bats (red); the black trace denotes head velocity. Eye-to-head velocity ratios (gains) in bats were ≤0.1 across all frequencies tested (**B**) Optokinetic reflex (OKR) gain plotted as a function of stimulus frequency for the same animals. The inset shows example eye-velocity traces at 0.2 Hz for mice (blue) and bats (red); the black trace denotes head velocity. In mice, OKR gain declined with increasing frequency, consistent with classical OKR–VOR crossover dynamics. In bats, OKR gain remained uniformly low (≤0.1) and relatively flat across the tested frequency range. (**C**) Micro-CT reconstructions of the skull and otic endocast of the right bony labyrinth in a mouse (top) and Seba’s short-tailed bat (bottom). Reconstructions are scaled and oriented to allow direct comparison of semicircular-canal morphology and planar orientation. Angles between ipsilateral canal pairs were similar across species, ranging from 89.8° (ASC–LSC) to 95.8° (LSC–PSC) in bats. In both species, the ASC–LSC angle was nearly orthogonal (±0.25–0.3° deviation). The ASC–PSC and LSC–PSC angles were obtuse, deviating from orthogonality by 4.9–12.9° in mice and 4.1–5.8° in bats. The ASC–PSC angle showed the largest species difference, being more obtuse in mice (102.9°) than in bats (94.1°). In both species, the LSC tilted upward anteriorly relative to the horizontal reference plane (+31.3° in mice; +44.8° in bats). See also Figure S2 and Table S1.

Together, these findings show that eye movement responses in *C. perspicillata* do not exhibit the frequency-dependent modulation or multisensory enhancement observed in mice. This pattern stands in stark contrast to the canonical mammalian gaze stabilization strategy exhibited by mice.

### Semicircular Canal Morphology Does Not Explain the Weak VOR

Finally, we compared semicircular canal morphology between bats and mice using high-resolution micro-CT imaging (Figure 4C). Measurements of canal radius of curvature and cross-sectional shape revealed only minor species differences. Notably, bats exhibited slightly more circular canal cross-sections, a geometry predicted to increase canal sensitivity based on established biophysical models. Detailed morphometric parameters for both species, including canal-plane angles, eccentricity values, and lumen radius measurements, are provided in Table S1. Together, these anatomical measurements indicate that the weak VOR observed in *C. perspicillata* cannot be readily explained by gross peripheral canal morphology.

## Discussion

For more than eight decades, the prevailing view in sensory physiology has been that bats do not move their eyes. This view originates from Walls’s influential book, the Vertebrate Eye^2^, in which he asserted that small nocturnal mammals—including bats—possess eyes that “never move even reflexively” (p.310). This categorical claim, derived entirely from anatomical inference rather than physiological measurement, became deeply embedded in the literature, despite never being experimentally tested. Our results correct this assertion by demonstrating robust reflexive eye movements in *Carollia perspicillata* and provide the first direct physiological evidence that bats possess an active oculomotor system. Specifically, Seba’s short-tailed bats generate strong, well-structured optokinetic eye movements with clear slow phases and compensatory quick phases, indicative of effective gaze stabilization. Notably, these robust responses occur despite a more limited oculomotor range (∼±10°) than that measured in mice (∼±20°) under the same stimulation.

Our empirical data aligns with a substantial but underappreciated body of neuroanatomical and behavioral evidence. Several echolocating bat species exhibit strong retinal projections to the dorsal lateral geniculate nucleus and superior colliculus, ^5,6^ and visual cues contribute to behaviors including homing, ^7–9^ escape responses, ^10–12^ obstacle avoidance, ^13,14^ and navigation. ^15^ In the absence of physiological measurements of eye movements, however, these observations remained difficult to reconcile with the claim that bat eyes are static. By directly quantifying gaze stabilization in bats, our study fills a longstanding gap in comparative research on eye movement control.

### Visual system characteristics and sensory integration in *Carollia perspicillata*

The sensory ecology of *Carollia perspicillata* suggests a potential function of eye movements in this species. This fruit bat inhabits dense tropical forests of Central and South America, where acoustic shadowing from foliage limits the range and precision of echolocation^16^. Although *C. perspicillata* can rely on echolocation alone for navigation and foraging, ^17,18^ it also uses vision to steer around obstacles and guide near-field behaviors. ^13^ Its moderately sized eyes, substantial binocular overlap, and rod-dominated retina that retains cone photoreceptors support visual function under dim light. ^19,20^ In addition, several bat species exhibit an area centralis or visual streak, indicating selective enhancement of spatial acuity or motion sensitivity. ^21,22^ In this ecological context, eye movements support visual information gathering and image stabilization, complementing acoustic sensing.

More broadly, bats exhibit marked diversity in visual system organization and in the relative weighting of vision and echolocation. Many insectivorous species that hunt fast-moving prey in complete darkness rely predominantly on echolocation and possess smaller eyes, along with enlarged auditory structures such as the inferior colliculus. ^23^ In contrast, visually guided species, including many fruit bats, show larger eyes and a more developed superior colliculus, consistent with robust visual function. ^23^ *C. perspicillata* exemplifies an intermediate sensory strategy, producing low-intensity, broadband echolocation calls with a limited operating range while relying more heavily on visual cues for navigation. These comparisons indicate that reliance on vision versus echolocation exists along a continuum shaped by ecological and behavioral demands rather than a strict sensory modality trade-off.

### A distinct sensory weighting strategy for gaze stabilization

A striking feature of our findings is the prominence of visual and otolith-driven contributions to gaze stabilization relative to semicircular canal–driven angular vestibulo-ocular reflexes. In most mammals, optokinetic and canal-driven vestibulo-ocular reflexes act in complementary fashion to stabilize gaze across stimulus frequencies. ^1^ In *C. perspicillata*, however, stabilization under passive conditions appears to be driven more heavily by visual and gravito-inertial cues, which departs from the canonical mammalian pattern.

These specializations appear relevant to the biomechanical demands of flight, but their contributions must be considered in the context of integrated vestibular processing. During natural flight, bats experience substantial translational, rotational, and gravitational forces associated with wingbeat oscillations and maneuvering. Maintaining alignment with the Earth-vertical axis therefore requires combined input from both otolith and semicircular canal systems. Otolith organs, which encode linear acceleration and head orientation relative to gravity, provide information relevant for head stabilization, while canal signals are essential for distinguishing tilt from translation. Consistent with this integrated control, bats preferentially stabilize their heads along the Earth-vertical axis during flight,^22^ and studies in freely flying Egyptian fruit bats show that such stabilization is achieved through coordinated modulation of wing and body kinematics that maintain a stable sensory scene.^23–25^ Further, evidence that echolocating horseshoe bats can perform Doppler shift compensation in the absence of acoustic feedback suggests that vestibular signals contribute to adaptive sensorimotor adjustments during flight.^26^ How vestibular signals are dynamically integrated and weighted across otolith and canal pathways during natural flight—particularly as a function of behavioral state—remains to be determined.

The reduced canal-driven angular vestibulo-ocular reflex observed under our under passive rotations conditions cannot be explained by peripheral anatomy. Across mammals, semicircular canal radius and geometry correlate with vestibular sensitivity, ^29,30^ and deviations from circularity reduce canal output. ^31–33^ *C. perspicillata* exhibits semicircular canal geometry comparable to that of mice, a morphology expected to favor high sensitivity. To date, there are no reports of peripheral hair-cell specializations or reductions in *C. perspicillata* comparable to those described in some aquatic species. ^34,35^ Together, these observations point to central mechanisms shaping vestibular function rather than peripheral constraints.

### Central gating and behavioral state dependence

We posit that canal-driven vestibulo-ocular pathways are selectively weighted with respect to behavioral state. In mammals, the vestibulo-ocular reflex is strongly modulated during voluntary gaze shifts and smooth pursuit to prevent interference with goal-directed movements^36–43^ as well as during voluntary eyelid closure. ^44^ State-dependent modulation of vestibular responses has also been demonstrated in animals with specialized locomotor modes, including pigeons during flight^45^ and tadpoles during swimming. ^46^

In bats, vestibulo-ocular modulation may be shaped by behaviors unique to flight and inverted posture. Wingbeat-induced accelerations, landing maneuvers, and roosting upside-down place a premium on gravito-inertial cues. Consistent with this idea, under microgravity conditions that disrupt otolith function, bats assume a neutral posture rather than the repetitive righting behaviors observed in other mammals, ^47^ suggesting functional vestibular adaptations. While distinguishing tilt from translation ultimately requires integration of both otolith and semicircular canal signals, ^48^ canal projections in bats may be preferentially routed toward non-ocular pathways, such as vestibulospinal or postural circuits, particularly during active flight or inverted roosting. ^49^ Such functional channeling could potentially account for the weak canal-driven angular vestibulo-ocular reflex observed under passive rotation conditions.

Together, our findings show that eye movements constitute an active component of gaze stabilization in bats, embedded within a sensory architecture that selectively reweights visual, otolith, and canal inputs. Rather than relying predominantly on semicircular canal–driven vestibulo-ocular reflexes, C. perspicillata emphasizes visual and otolith cues under passive conditions while likely engaging canal pathways during active flight. Through empirical demonstration of robust reflexive eye movements in *C. perspicillata*, our study reshapes understanding of bat sensorimotor control and provides a framework for comparative investigations of gaze stabilization across species with diverse locomotor and sensory specializations.

## Resource availability

### Lead contact

Request for further information and resources should be directed to and will be fulfilled by the lead contact, Kathleen E. Cullen.

### Materials availability

This study did not generate new, unique reagents

### Data and code availability

- Data have been deposited at figshare and are publicly available at https://doi.org/10.6084/m9.figshare.31428821
- This paper does not report original code.
- Any additional information required to reanalyze the data reported in this paper is available from the lead contact upon request.

## Acknowledgements

We thank Chenhao Bao for his valuable contributions to signal processing and quantitative analysis. We also gratefully acknowledge Te K. Jones for assistance with animal procedures and for invaluable support in troubleshooting experimental setups and protocols. This work was supported by the National Institutes of Health grants 1U01-NS111695 (KEC), NIH R01 NS121413 (CFM), Human Frontiers Science Program Research Grant RGP0045/2022 (CFM), NSF CRCNS Grant 2011619 (CFM), and Office of Naval Research Grants N00014-23-1-2086 and N00014-17-1-2736 (CFM).

## Author Contributions

K.E.C., H.H.V.C., C.F.M, and G.C., designed the study.

H.H.V.C., T.J, D.S, and G.C. performed the experiments. H.H.V.C. and G.C. analyzed the data.

H.H.V.C. and G.C. prepared the figures and tables. H.H.V.C. and K.E.C. wrote the paper with input from C.F.M, and G.C.

## STAR Methods

### Experimental model details

#### Animals

Four captive-born Seba’s short-tailed bats used in this study were group-housed in a colony room and maintained at 24-28°C and 40-70% relative humidity. Bats had access to sufficient roosting locations, room to fly, and were provided ad libitum access to fruit, nectar, and water. In addition, six mice were obtained from The Jackson Laboratory and maintained on C57BL/6J background. Mice were group-housed with their littermates on a 12:12 h light: dark cycle at 20°C with ad libitum access to food and water. All animal procedures were approved by the Institutional Animal Care and Use Committees at Johns Hopkins University.

### Method details

#### Experimental setup

Animals were positioned in a padded body tube to minimize body movements. The implanted headpost was rigidly fixed to a head-fixation bar attached to the body tube. The assembly was placed on a motorized turntable surrounded by a visual surround composed of vertical black-and-white stripes (visual angle width of 5°).

For constant-velocity VOR and OKR testing, either the turntable (for VOR) or the visual surround (for OKR) was rotated clockwise and counterclockwise at 50°/s for 72 s. The platform accelerated from 0 to 50°/s within 500 ms and maintained at the constant velocity for 72 s (10 full rotations). VOR measurements were performed in darkness, and OKR measurements under illuminated conditions. Each constant-velocity OKR and VOR condition was repeated 3–5 times per animal. Slow-phase eye velocity during constant-velocity stimulation was quantified for each animal and summarized at the group level across animals.

For angular VOR and OKR testing, the turntable or visual surround was rotated sinusoidally at 0.2, 0.4, 0.8, 1, and 2 Hz with peak velocities of ±16°/s. VOR responses were recorded under both dark and light conditions. Gain and phase were computed from five consecutive steady-state cycles at each stimulus frequency.

For OVAR measurements followed previously published procedures. ^50^ Briefly, the motorized turntable tilted 17° relatively to the horizontal. The turntable accelerated from 0 to 50°/s within 500 ms and maintained at the constant velocity for 72 s (10 full rotations). Eye movement data were collected in bats and mice using an infrared video system (ETL-200, ISCAN system) mounted on the turntable. Head velocity of the animal was simultaneously measured using a MEMS sensor (MPU-9250, SparkFun Electronics). OVAR stimuli were repeated 3–5 times per animal. Slow-phase eye velocity during OVAR stimulation was quantified and summarized at the group level across animals.

#### CT measurements

Mice (n=3) and bats (n=3) were scanned using a Bruker SkyScan 1172 micro-CT scanner (imaging parameters are provided in Table S2). Image stacks were digitally segmented using 3D Slicer (v. 4.11.20210226)^51^ to generate three-dimensional reconstructions of the cranial bones and the otic endocast. From these reconstructions, we measured linear dimensions of the semicircular canals, including height (maximum distance from the vestibule to the center of the canal lumen), width (maximum distance between the two arms of the canal, perpendicular to the height), and radius of the canal lumen for the left and right ears of each specimen.

We calculated the radius of curvature (R) of each canal as

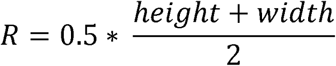

following previously published methods.^52^ To assess the ellipticity of each semicircular canal, we calculated canal eccentricity (*ecc*) using the equation

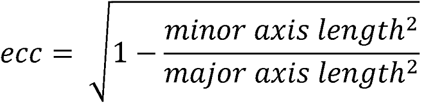

as previously described,^32^ where values closer to 1 indicate greater ellipticity and values closer to 0 indicate greater circularity.

We measured streamline lengths of each semicircular canal by extracting midline curves along the centroid of the lumen of each canal using the VMTK extension in 3D Slicer.^53^ The planar relationships of ipsilateral semicircular canal pairs were characterized by fitting planes along canal midlines and measuring the angle between the planes of two canals in 3D Slicer. We additionally calculated angular deviation of the lateral semicircular canal midline from a horizontal reference plane defined by landmarks at the center points of the external auditory canals and the inferior-most margin of the orbits, representing a stereotaxic horizontal reference used in previous studies of semicircular canal orientation.^54–56^

### Quantification and statistical analysis

Eye position signals were acquired digitally at 240 Hz using an infrared video-based eye-tracking system (ISCAN). Head velocity was acquired at 1 kHz using a MEMS inertial sensor. Both signals were low-pass filtered offline at 125 Hz prior to further analysis. Eye position signals were then numerically differentiated to obtain eye velocity. For all analyses, pre-and post-stimulus periods were carefully examined to confirm that eye movements emerged only following stimulus onset. Specifically, we verified that the 95% confidence interval of the baseline (pre-stimulus) eye velocity did not overlap with that of the post-stimulus response, ensuring that the measured eye movements were stimulus-evoked rather than spontaneous fluctuations. Quick phases were detected using a velocity threshold algorithm (75°/s)^57,58^ and excluded from further analysis, leaving only uninterrupted slow-phase segments; no interpolation across quick phases was performed. For angular VOR and OKR, gains and phases were computed from five consecutive sinusoidal cycles for each frequency.

VOR and OKR gains and phases were estimated by least-squares optimization. For OVAR, the time constant of the slow-phase eye-velocity decay as well as the frequency and bias of resulting sinusoidal modulation were determined. Maximum ocular deviation was defined as the largest slow-phase eye-position displacement from baseline during the stimulus. Data are presented as mean ± SEM across all tested frequencies for mice and bats, unless otherwise indicated.

