## Supplemental Figure 1-2, Supplemental Table 1-2 for "Bat eye movements resolve a long-standing question in gaze control"

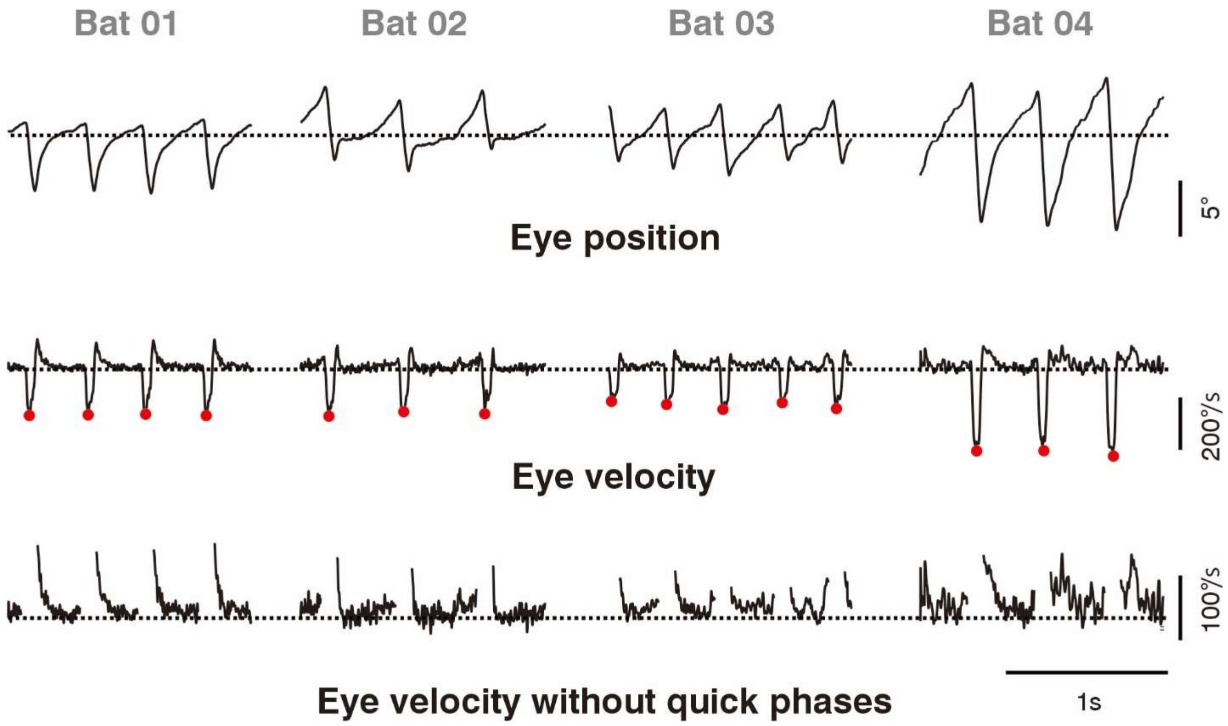


**Figure S1. Two-component structure of optokinetic slow phases is consistent across individual bats. Related to Figure 1.** Representative eye position (top), eye velocity (middle), and slow-phase eye velocity with compensatory quick phases removed (bottom) are shown for four individual Seba's short-tailed bats (Bat 01–Bat 04) during optokinetic stimulation. Traces illustrate repeated cycles of optokinetic responses, with quick phases identified by red markers in the eye-velocity traces. Across animals, individual slow phases consistently exhibit two components, both in the direction of the visual stimulus: an initial, brief higher-velocity transient followed by a slower tracking component. The timing and magnitude of this initial transient vary across cycles and individuals, contributing to the irregular appearance of the population-averaged slow-phase velocity traces shown in Figure 1C–D. Scale bars indicate eye position (5°), eye velocity (200°/s), slow-phase velocity (100°/s), and time (1 s).

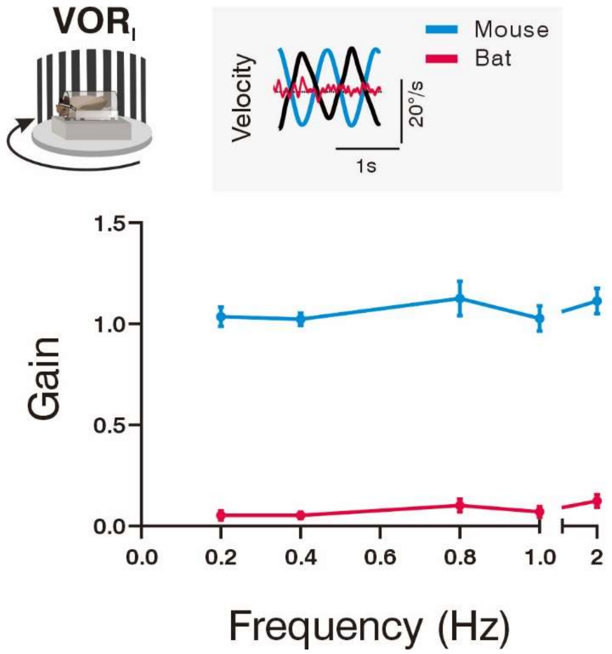


**Figure S2. Seba's short-tailed bat exhibits minimal to no compensatory response during the angular vestibulo-ocular reflex in light. Related to Figure 4.** VOR$_l$ gain (mean ± SEM) is plotted as a function of stimulus frequency for control mice (blue) and Seba's short-tailed bats (red). The inset shows example eye-velocity traces at 0.8 Hz for mice (blue) and bats (red); the black trace denotes head velocity. Animals were rotated sinusoidally about the Earth-vertical axis while the patterned visual surround remained stationary. In this paradigm, both visual and vestibular cues are available, allowing the angular VOR and OKR to operate synergistically to stabilize gaze. As expected, mice generated robust compensatory eye movements across all tested frequencies, with gains approaching 1. In contrast, bats produced minimal to no compensatory eye movement under these combined-cue conditions, resulting in stabilization gains near 0.

| | | Mouse (n=3) | Seba's short-tailed bat (n=3) |
|---|---|---|---|
| **ASC** | Posterior braincase width (mm) | 10.43 ± 0.61 | 11.14 ± 0.09 |
| | Height (mm) | 1.30 ± 0.05 | 1.52 ± 0.06 |
| | Width (mm) | 1.92 ± 0.06 | 1.62 ± 0.10 |
| | Length (mm) | 3.66 ± 0.09 | 3.13 ± 0.09 |
| | Lumen radius (mm) | 0.08 ± 0.01 | 0.07 ± 0.00 |
| | Radius of curvature (mm) | 0.80 ± 0.03 | 0.78 ± 0.04 |
| | Eccentricity | 0.74 ± 0.01 | 0.34 ± 0.05 |
| **LSC** | Height (mm) | 1.03 ± 0.01 | 1.25 ± 0.02 |
| | Width (mm) | 1.42 ± 0.03 | 1.50 ± 0.06 |
| | Length (mm) | 2.91 ± 0.03 | 3.35 ± 0.15 |
| | Lumen radius (mm) | 0.08 ± 0.00 | 0.08 ± 0.00 |
| | Radius of curvature (mm) | 0.61 ± 0.01 | 0.69 ± 0.02 |
| | Eccentricity | 0.69 ± 0.02 | 0.55 ± 0.03 |
| **PSC** | Height (mm) | 1.52 ± 0.03 | 1.35 ± 0.12 |
| | Width (mm) | 1.18 ± 0.03 | 1.31 ± 0.11 |
| | Length (mm) | 3.24 ± 0.15 | 3.20 ± 0.06 |
| | Lumen radius (mm) | 0.08 ± 0.01 | 0.08 ± 0.00 |
| | Radius of curvature (mm) | 0.68 ± 0.01 | 0.66 ± 0.06 |
| | Eccentricity | 0.63 ± 0.02 | 0.25 ± 0.06 |
| **Canal angle relationships** | ASC ∡ LSC | 89.75° ± 0.84° | 89.70° ± 0.99° |
| | ASC ∡ PSC | 102.93° ± 0.72° | 94.13° ± 0.81° |
| | LSC ∡ PSC | 94.90° ± 1.65° | 95.78° ± 0.43° |
| | LSC ∡ Horizontal reference | 31.30° ± 0.89° | 44.77° ± 1.78° |

**Table S1. Summary of semicircular canal measurements. Related to Figure 4.**
Values represent species means ± standard deviation.

| Species | ID | kV | µA | Projections | Filter | Frame Averaging | Voxel size |
|---|---|---|---|---|---|---|---|
| ***Mus musculus*** | MM1 | 100 | 100 | 785 | Al 0.5 mm | 10 | 13.53 um |
| | MM2 | 55 | 181 | 2331 | Al 0.5 mm | 2 | 9.91 um |
| | MM3 | 85 | 118 | 1159 | Al 0.5 mm | 10 | 13.53 um |
| ***Carollia perspicillata*** | CP1 | 100 | 100 | 962 | Al 0.5 mm | 10 | 13.53 um |
| | CP2 | 55 | 181 | 2309 | Al 0.5 mm | 2 | 9.91 um |
| | CP3 | 100 | 100 | 1125 | Al 0.5 mm | 12 | 13.53 um |

**Table S2. Micro-CT scanning parameters. Related to STAR Methods.**
